# The bacillithiol pathway is required for biofilm formation in *Staphylococcus aureus*

**DOI:** 10.1101/496653

**Authors:** Megha Gulati, Jason Thomas, Mamta Rawat, Clarissa J. Nobile

## Abstract

*Staphylococcus aureus* is a major human pathogen that can cause infections that range from superficial skin and mucosal infections to life threatening disseminated infections. *S. aureus* can attach to medical devices and host tissues and form biofilms that allow the bacteria to evade the host immune system and provide protection from antimicrobial agents. To counter host-generated oxidative and nitrosative stress mechanisms that are part of the normal host responses to invading pathogens, *S. aureus* utilizes low molecular weight (LMW) thiols, such as bacillithiol (BSH). Additionally, *S. aureus* synthesizes its own nitric oxide (NO), which combined with its downstream metabolites may also protect the bacteria against specific host responses. We have previously shown that LMW thiols are required for biofilm formation in *Mycobacterium smegmatis* and *Pseudomonas aeruginosa*. Our data show that the *bshC* mutant, which is defective in the last step of the bacillithiol pathway and lacks BSH, is impaired in biofilm formation. We also identify a putative *S*-nitrosobacillithiol reductase (BSNOR), similar to a *S*-nitrosomycothiol reductase found in *M. smegmatis*, and show that the BSNOR mutant has reduced levels of BSH and decreased biofilm formation. Our studies also show that NO plays an important role in biofilm formation and that acidified sodium nitrite severely reduces biofilm thickness. These studies provide insight into the roles of oxidative and nitrosative stress mechanisms on biofilm formation and indicate that bacillithiol and nitric oxide are key players in normal biofilm formation in *S. aureus*.

**Importance:** *Staphylococcus aureus* is the most frequent cause of biofilm-associated infections in hospital settings. The biofilm mode of growth allows the pathogen to escape the host immune response and is extremely difficult to combat, as biofilms are highly resistant to physical and chemical stressors. As outbreaks of antibiotic-resistant bacterial strains become more commonplace, it is essential to understand the pathways involved in biofilm formation in order to target this important virulence factor. Low molecular weight thiols enable *S. aureus* to combat oxidative and nitrosative stress mechanisms that are used by host cells to defend against infection. Our findings indicate that bacillithiol and nitric oxide, which are produced by *S. aureus* to combat these host generated stressors, are important for biofilm development, and that disruption of these pathways results in biofilm defects. In the long term, this work may lead to new solutions to eradicate *S. aureus* biofilms in the clinic.

## Introduction

***S****taphylococcus aureus* is a gram-positive, spherical-shaped bacteria that is a major human pathogen capable of causing both superficial as well as life-threatening systemic and chronic infections in humans (1, 2). *S. aureus* is also a commensal microorganism that can reside on the skin, and within the nostrils, throat, perineum and axillae of 25-30% of healthy humans (3-6). One *S. aureus* virulence factor that is particularly challenging to manage in hospital settings is its ability to form biofilms, surface-attached communities of cells that are encased in an extracellular polymeric matrix (7). These biofilms can form on both biotic and abiotic surfaces and are more resistant to antimicrobial agents and the host immune response compared to their free-floating (planktonic) counterparts (7-10). *S. aureus* was one of the first microbial species found to grow as a biofilm on medical devices in clinical settings and is responsible for frequent episodes of bacteremia and sepsis in hospital settings (11). In fact, *S. aureus* biofilms, including those formed by methicillin-resistant *S. aureus* (MRSA) strains, are the leading cause of hospital-acquired sepsis, commonly observed in patients with burn wounds and implanted or indwelling medical devices (12-16).

The host immune system employs various mechanisms to recognize and counter infections by *S. aureus* and other invading pathogens (17-19). Neutrophils and macrophages are among the first lines of defense against epithelial invasion by *S. aureus*, leading to a release of reactive oxygen species (ROS) to combat *S. aureus* colonization (20-22). In particular, activated neutrophils mount an oxidative burst, where NADPH oxidase produces ROS, such as superoxide (O_2_^-^) and hydroxyl radical (·OH) (23, 24). Consistent with this protective oxidative burst, mouse models of chronic granulomatous disease that show a severe reduction in NADPH oxidase are highly susceptible to infections by *S. aureus* (25, 26). *S. aureus* proteins are particularly vulnerable to damage by ROS, where ROS oxidize the thiol residues in cysteine moieties (18). Neutrophils also release cytokines and chemokines, which in turn can activate macrophages at the site of *S. aureus* infection. Activated macrophages release high concentrations of nitric oxide (NO), which can combine with superoxide to form highly toxic anions, such as peroxynitrite (OONO^-^) (27). These high concentrations of NO and other reactive nitrogen species (RNS), which are formed by reactions of NO with other oxidants, together produce nitrosative stress that is useful in eliminating invading bacterial cells (24). To counter these oxidative and nitrosative host stress mechanisms, *S. aureus* in turn has evolved several pathogen-derived adaptive mechanisms to survive and propagate within the host (20, 28-31).

One pathogen-derived mechanism is the production of low molecular weight (LMW) thiols that are thought to be crucial in providing protection to cytosolic proteins against ROS, other reactive electrophilic species, antibiotics, and heavy metals (32). LMW thiols oxidize more slowly than cysteine and can play crucial roles in redox reactions and maintaining redox homeostasis (33). Glutathione (GSH), a tripeptide, is the most abundant LMW thiol produced in cells of eukaryotes and many gram-negative bacteria (34, 35); most gram-positive bacteria, however, are thought to lack GSH (35). Another LMW thiol, mycothiol (MSH), was identified as the primary thiol in high G+C gram-positive bacteria (36, 37), while bacillithiol (BSH) was identified as the primary LMW thiol in the low G+C firmicutes (38). MSH and BSH differ considerably in structure from GSH by containing an amino sugar glucosamine backbone instead of a peptide backbone. BSH biosynthesis and function has been elucidated in the firmicutes *S. aureus* and *B. subtilis* (38), and *S. aureus* mutants lacking BSH have been shown to display increased sensitivities and susceptibilities to killing by oxidative stress (39, 40). One study demonstrated that BSH can reduce oxidants such as H_2_O_2_ directly and that BSH may participate in a general electron relay with bacilliredoxins to reduce oxidants (41).

LMW thiols have also been implicated in protection against nitrosative stress (42). NO can react with GSH to form GSNO spontaneously, preventing its reaction with cysteine moieties (43). The amount of GSNO in the cell is modulated by *S*-nitrosoglutathione reductase (GSNOR), which regenerates GSH and reduces NO to ammonia, and also has a dual function as a *S*- (hydroxylmethyl) glutathione dehydrogenase, participating in formaldehyde detoxification (44). The role of GSNOR in virulence and pathogenesis has been elucidated in *Streptococcus pneumoniae*, where GSNOR is required for survival in blood (44). Paradoxically, *S. aureus* synthesizes its own NO using a bacterial nitric oxide synthase (SaNOS), which suggests that NO may also serve some protective role for the bacterium (45). NO and its downstream metabolite, such as nitrite, may protect *S. aureus* and other NOS-producing bacteria by scavenging HClO to produce nitrate, which is less reactive (18). NO can also block cysteine oxidation by *S*-nitrosylating exposed cysteines (46) and activating catalases that reduce the concentration of harmful H_2_O_2_ (18). Overall, SaNOS mutants display increased endogenous ROS and superoxide levels (47). Recent studies also find that the production of NO by *S. aureus* is essential for nasal colonization and skin abscess development, and mutants lacking the enzyme show decreased virulence (48-50). Interestingly, at low concentrations NO can act as a signaling molecule in a dose-dependent manner to modulate group activities, such as biofilm formation in numerous bacterial species (51). These complex and multifactorial roles for NO at different concentrations in bacteria are analogous to that in mammalian cells (51).

In *S. aureus*, one report found that NO-mediated signal transduction regulates biofilm formation and dispersal, although the mechanistic details of the regulation are unknown (52). NO and nitrite (NO_2_^-^) impair polysaccharide intercellular adhesion (PIA)-dependent biofilm formation in *S. aureus* (53); PIA is a major constituent of the extracellular matrix of staphylococcal biofilms. It was also recently shown that NO inhibits *S. aureus* virulence by disrupting intercellular communication between bacterial cells by targeting proteins involved in quorum sensing (46). Biofilm formation in *P. aeruginosa* is also affected by NO in a concentration dependent manner and the bacteria harbor NO-responsive regulators that modulate biofilm dispersal (54, 55).

We previously reported that MSH and a *S*-nitrosomycothiol reductase are required for biofilm formation in *Mycobacterium smegmatis* (56). We also recently reported that a *P. aeruginosa* mutant lacking GSH is defective in biofilm formation (57). These studies indicate that regardless of the structure of the LMW thiols, these thiols play important roles in biofilm formation in diverse bacterial species. Here, we report a new role for BSH in *S. aureus* biofilms. We show that like GSH and MSH mutants, BSH mutants are also impaired in biofilm formation. We confirm that NO is involved in biofilm formation and identify a putative *S*-nitrosobacillithiol reductase in *S. aureus*. Overall, both ROS and RNS stressors are encountered by *S. aureus* biofilms (23), however the detailed mechanisms by which biofilms counter these stressors and how thiols play a role in response to them are unknown.

## Results

### *S*. *aureus* contains a putative *S*-nitrosobacillithiol reductase

Recently, we described the characterization of a *M. smegmatis* mutant disrupted in *S*-nitrosomycothiol reductase (MSMEG_4340) (56). This mutant along with mutants lacking the LMW thiols, MSH and ergothioneine (ESH), was more susceptible to *S*-nitrosoglutathione (GSNO) (56). Since MSH is structurally similar to BSH, we reasoned that the *S*-nitrosobacillithiol reductase (BSNOR) would be similar in sequence to the mycobacterial protein. MSMEG_4340 amino acid sequence and BLASTp analysis was used to identify the protein with the most amino acid identity/similarity in *S. aureus* (strain SAUSA300_FPR375). ORF SAUSA300_0055 contained the most identity at 35% (127/357) and similarity at 54% (193/357) with 96% coverage to MSMEG_4340 (E-value = 1e^-72^). We obtained a *S. aureus* USA300_FPR375 LAC JE2 transposon mutant disrupted in this ORF from the “Network on Antimicrobial Resistance in *Staphylococcus aureus*” through BEI Resources to characterize its role in biofilm formation and response to oxidative and nitrosative stressors. Under standard growth conditions in rich TSB media, the mutant grew similar to wildtype (WT). Surprisingly, the mutant also grew similarly to WT when treated with GSNO and sodium nitrite (Supplemental Figure 1). Since *S*-nitrosothiol reductases have a dual activity as formaldehyde dehydrogenases (58, 59), growth on formaldehyde was also assessed, and we found that there was no difference in growth from WT (Supplemental Figure 1).

### MIC for Diamide is higher under biofilm conditions compared with planktonic growth

We determined the MIC for diamide, a thiol oxidant, DTT, a reductant, and sodium nitrite, which releases NO chemically in an acidic solution at pH less than 7, to cause nitrosative stress (60) for the wildtype (WT) *S. aureus* strain under both planktonic and biofilm conditions. The MIC was 3.91 mM, 250 mM, and 31.25 mM for diamide, DTT, and sodium nitrite, respectively for WT grown under planktonic culture conditions (Table 1). The MIC for DTT and sodium nitrite was the same for the WT grown under biofilm conditions (Table 1). There was an approximate two-fold increase in the WT MIC for diamide under biofilm conditions, indicating that biofilms are more resistant to this oxidant.

**Table 1.**

|  | MIC Planktonic |  |  | MIC Biofilm |  |  |
| --- | --- | --- | --- | --- | --- | --- |
| Strain | DTT | Diamide | Sodium Nitrite | DTT | Diamide | Sodium Nitrite |
| JE2 | 250 mM | 3.91 μM | 31.25 mM | 250 mM | 7.81 mM | 31.25 mM |
| BshC | 250 mM | 3.91 μM | 31.25 mM | 250 mM | 7.81 mM | 62.50 mM |
| BsnoR | 250 mM | 3.91 μM | 31.25 mM | 250 mM | 7.81 mM | 31.25 mM |

### Bacillithiol levels decrease in the BSNOR mutant upon treatment with sodium nitrite

Since BSH can buffer NO by forming BSNO, we measured BSH levels upon exposure to sodium nitrite in the WT and BSNOR transposon mutant. As the MIC for sodium nitrite was 31.25 mM, we chose 20 mM sodium nitrite as the sublethal concentration for testing the effect of sodium nitrite on BSH levels. Treatment with 20 mM sodium nitrite did not cause cell death as indicated by a lack of change in CFUs before or after 30 mins of sodium nitrite treatment in both WT and BSNOR mutant strains. However, treatment with sodium nitrite resulted in a decrease in BSH levels from 1.21 to 0.81 µmol/g dry weight in the WT and a decrease in BSH levels from 1.15 to 0.62 µmol/g dry weight in the BSNOR transposon mutant. There was no difference in BSH levels in the untreated WT and BSNOR mutant, but the BSNOR mutant had significantly less BSH levels relative to WT after exposure to sodium nitrite (Figure 1).

**Figure 1.**
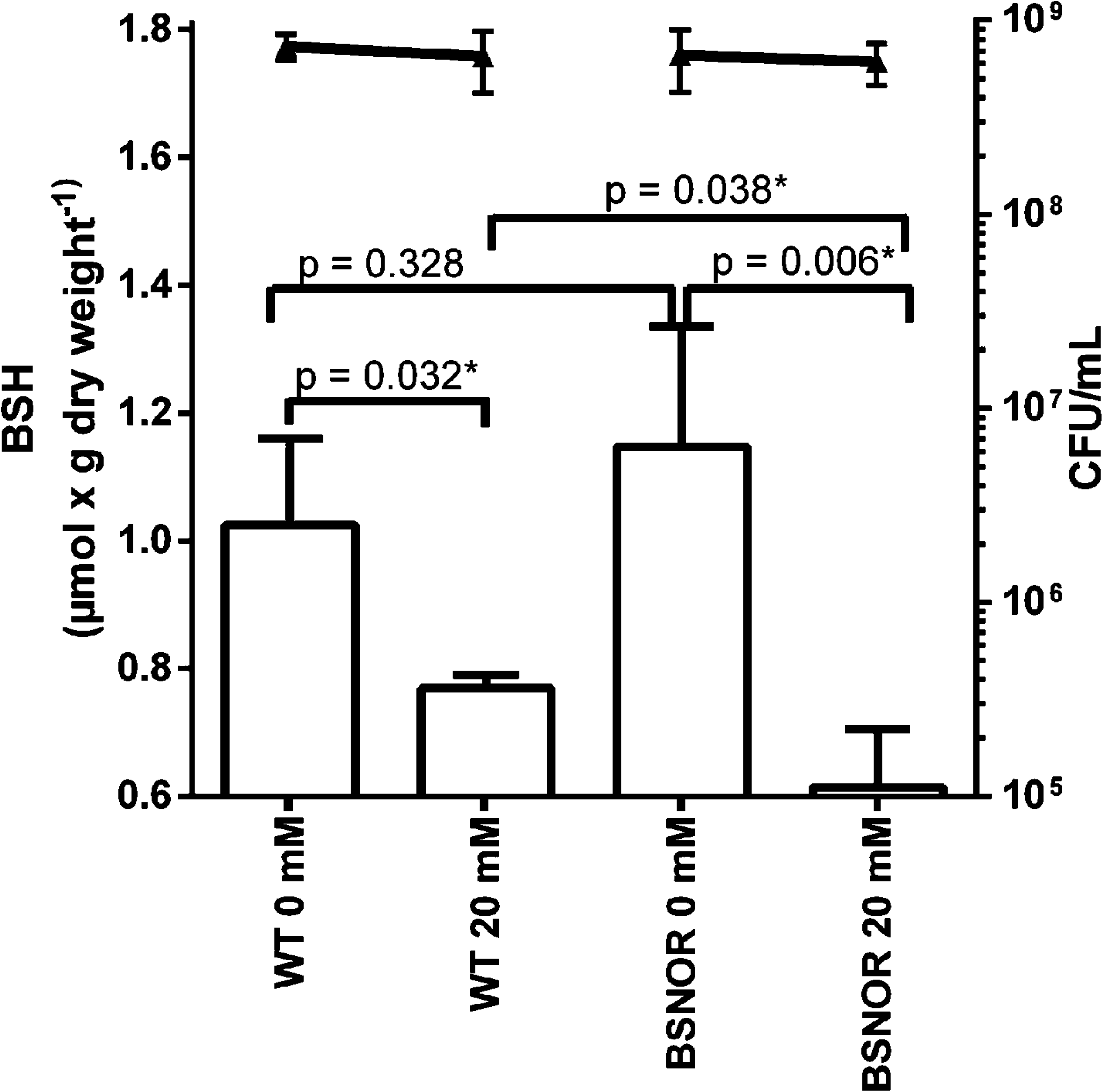
Reduced Bacillithiol level in the BSNOR mutant after treatment with sodium nitrite. BSH levels measured in *S. aureus* wild type (WT) cells and BSNOR mutant using HPLC analysis. BSH levels were measured in both cell types with no treatment and following treatment with sublethal concentrations of sodium nitrite (20 mM). The experiment was performed in quadruplicate and data presented represents mean and standard deviation. The right axis indicates the CFU/mL count which is represented by the triangle with error bars for standard deviation. The left axis indicates the bacillithiol concentrations that are represented by bars with standard deviation error bars. A student t-test was performed to determine significance; p-values ≤ 0.05 (*).

### Sodium nitrite has no effect on *bshC* and BSNOR expression

To determine if the decrease in BSH was due to a decrease in BSH biosynthesis, *bshC* expression levels were quantified using qPCR on WT cultures treated with sodium nitrite. At 5 min post-treatment, *bshC* is repressed (0.47) and remains repressed even after 30 min of sodium nitrite treatment. This data indicates that sodium nitrite causes repression of BSH biosynthesis through transcriptional control (Figure 2). Surprisingly, BSNOR expression does not change after 5 min or 30 min of sodium nitrite exposure. The positive control, the catalase *katA*, is induced three-fold (53).

**Figure 2.**
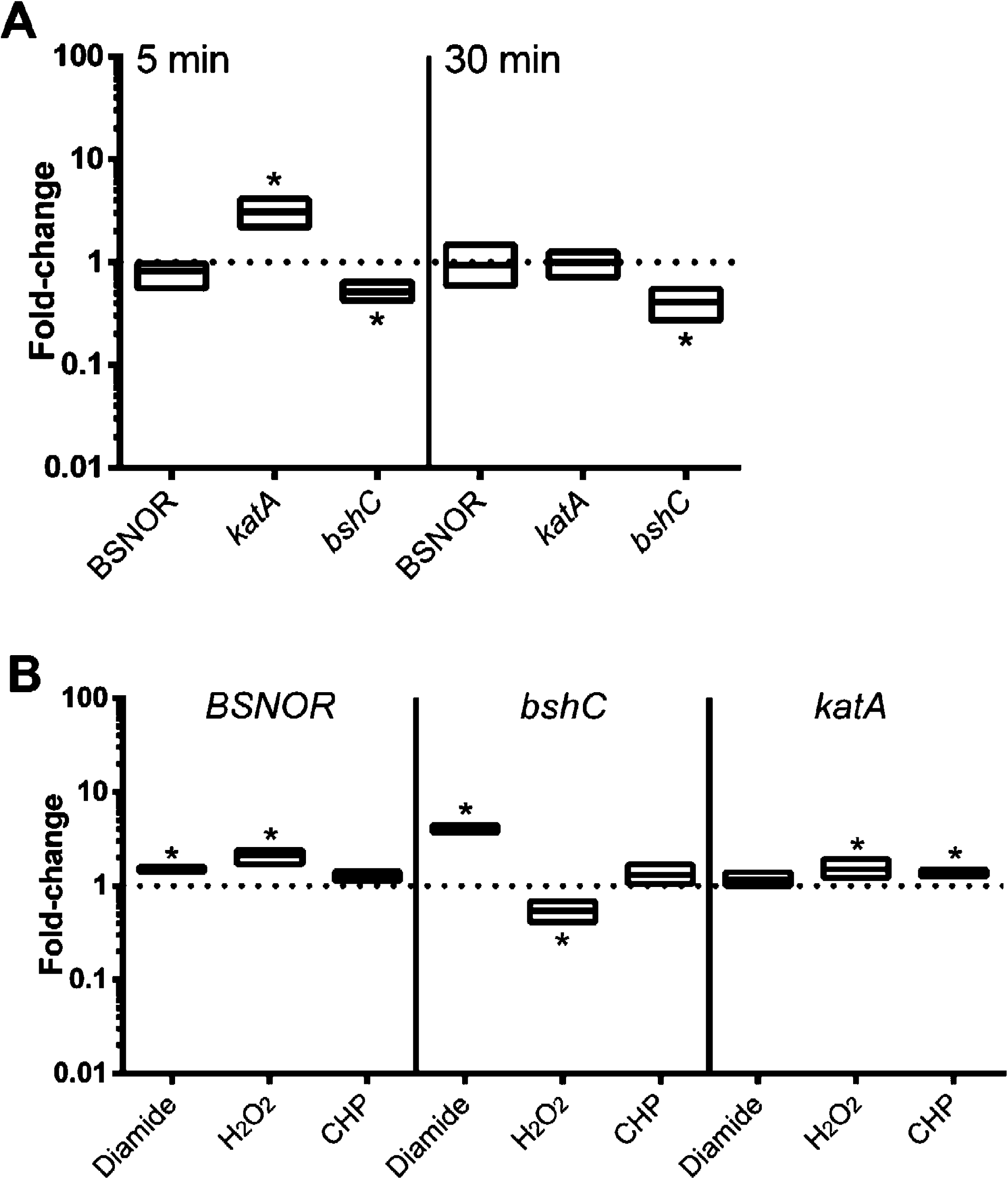
Expression levels of *BSNOR, bshC*, and *katA* genes under various stressors. A. Expression of *BSNOR, bshC*, and *katA* treated with 20 mM sodium nitrite for 5 min or 30 min in *S. aureus* wildtype. B. Expression of *BSNOR, katA* and *bshC* under oxidative stress (5 mM diamide, 100 µM cumene hydroperoxide, and 100 µM H_2_O_2_ for 30 min) in *S. aureus* wildtype. Relative gene expression was calculated using the 2^-ΔΔCT^ method (ΔCt = Ct sample - Ct control) and reported as fold change in gene expression of each sample normalized to the *gyrB* housekeeping gene relative to the untreated culture. A Mann-Whitney U test was performed to determine significance; p-values ≤ 0.05 (*).

We also checked *bshC* and BSNOR expression after treatment with oxidants (Figure 2). Upon exposure to hydrogen peroxide, BSNOR is induced two-fold while *katA* is only induced slightly (1.51) and *bshC* is repressed (0.55). Neither BSNOR nor *bshC* are affected by treatment with the lipid peroxide, cumene hydroperoxide, and *katA* is induced weakly (1.36). Treatment with the thiol oxidant, diamide, results in a slight upregulation of BSNOR (1.5), upregulation of *bshC* (4-fold), and no change in *katA* expression.

### BshC and BSNOR are needed for normal biofilm development

To determine if BshC and BSNOR are needed for biofilm formation, we tested the *bshC* and BSNOR transposon mutants and compared them with WT. Biofilms were allowed to develop for 24 hours, at 37°C on polystyrene plates and imaged using confocal scanning laser microscopy (CSLM). Our results show that either gene deletion results in a reduced biofilm depth (∼46 µm) compared to the wildtype biofilm (∼80 µm) (Figure 3).

**Figure 3.**
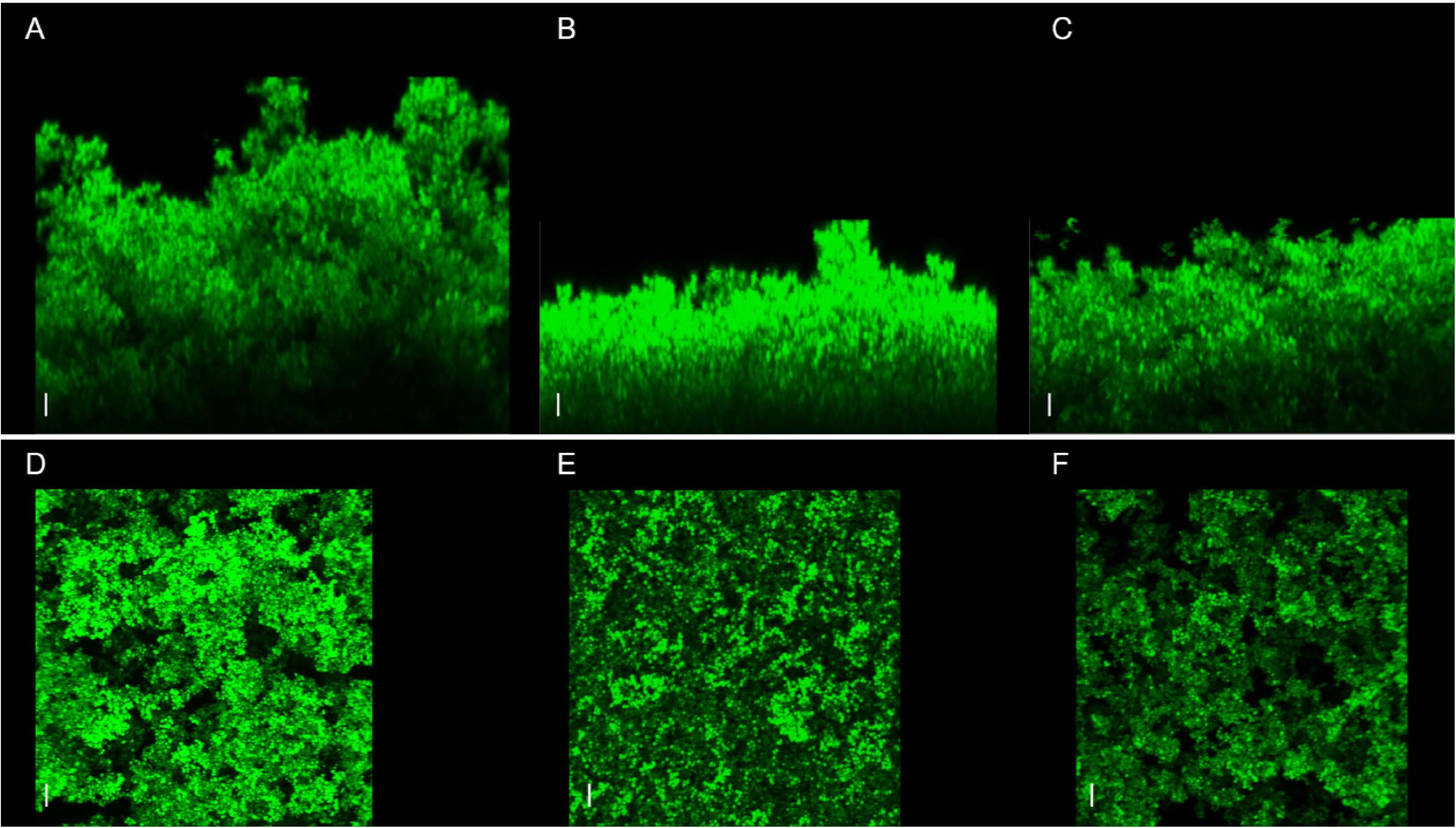
Confocal scanning laser microscopy (CSLM) images of *S*. *aureu*s 24 hr biofilms. Images show depth (side view) and top view of biofilms formed by wild-type JE2 (A and D), Δ*bshC* (B and E), and Δ*bsnoR* (C and F). Biofilms were formed on the bottom of 6-well polystyrene plates and stained with 2 μM Syto9 for 1 hr in the dark. Biofilms were imaged using an LSM-700 confocal microscope with a 63x objective and analyzed using ZEN software. Scale bars are shown in white in each panel and represent 5 µm.

### Diamide, DTT, and sodium nitrite reduce biofilm formation in *S*. *aureus*

To determine if oxidative, reductive, and nitrosative stressors can inhibit biofilm formation, we performed a biofilm MIC assay on polystyrene plates using a sustained inhibition biofilm assay (61), where the compound (diamide, DTT, or nitrite) is present during cell adherence and during the period of biofilm formation (24 hr). Our results show that 7.81 mM of diamide, 250 mM DTT, or 31.25 mM of sodium nitrite completely inhibits wild type biofilm formation in wildtype, Δ*bshC*, and Δ*bsnoR* strains (Table 1).

We also visualized the WT biofilm formed in the presence of sublethal concentrations of diamide, DTT and sodium nitrite by CSLM. Our results indicated that even sublethal concentrations of DTT (150 mM) severely reduced biofilm thickness to ∼ 20 µm (Figure 4B, 4E) and sublethal concentrations of diamide (1.95 mM) reduced biofilm thickness to ∼35 µm (Figure 4C, 4F). Since acidified TSB media was used for sodium nitrite treatments, we tested whether the decrease in biofilm thickness was due to the pH or sodium nitrite. Our results show that acidified TSB media reduces biofilm thickness to ∼45 µm compared with ∼80 µm biofilms formed in standard TSB media (Figure 4A, 4D and Figure 5A, 5C). However, sodium nitrite further reduced biofilm thickness to ∼10 µm in acidified TSB media (Figure 5B).

**Figure 4.**
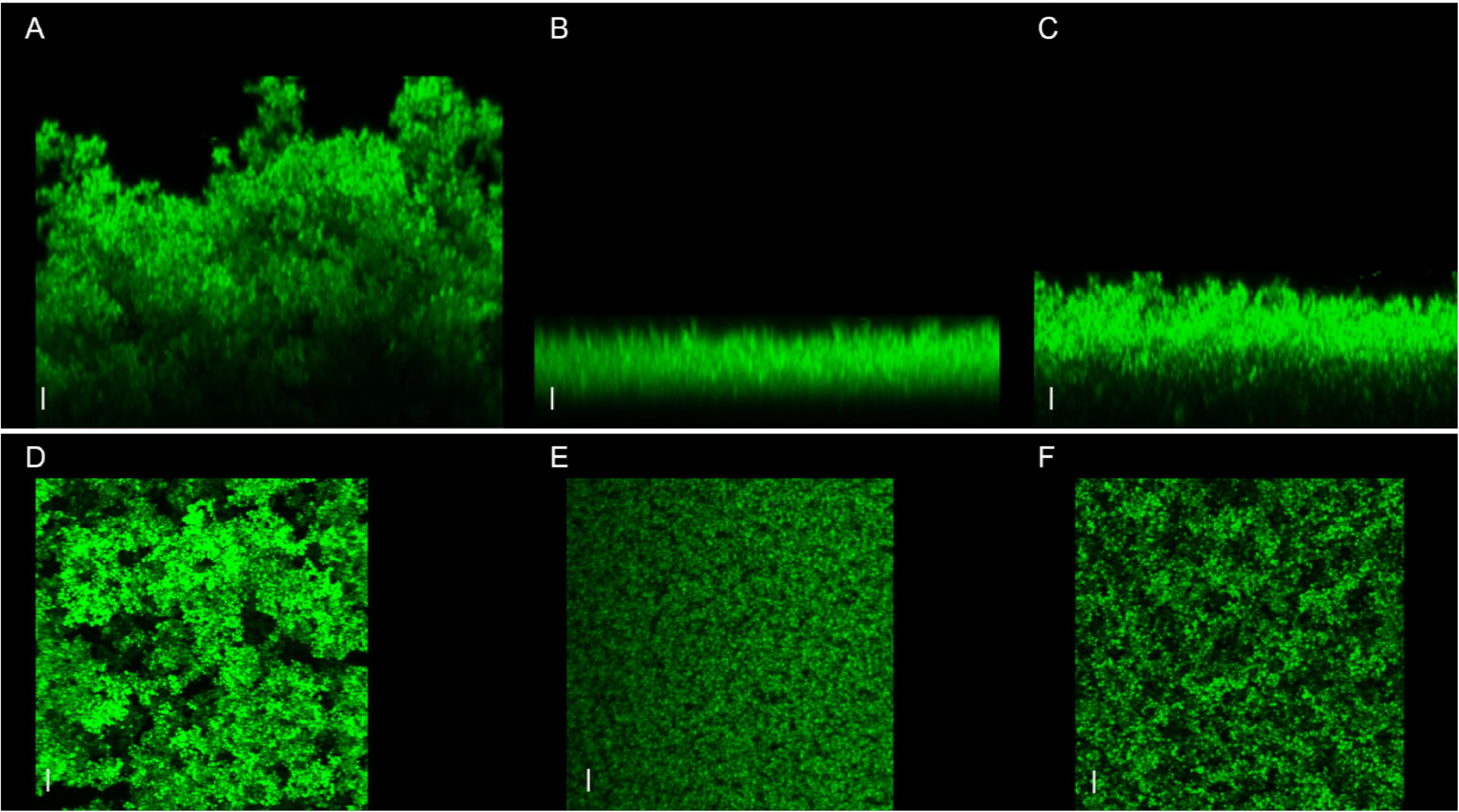
Effects of sublethal concentrations of DTT and diamide on *S*. *aureus* biofilms by CSLM. Images show depth (side view) and top view of biofilms formed by wild-type JE2 in TSB media supplemented with 0.75% glucose (TSB-G) (A and D), TSB-G media and 150mM DTT (B and E), and TSB-G media with 1.95 mM diamide (C and F). Biofilms were formed on the bottom of 6-well polystyrene plates and stained with 2 μM Syto9 for 1 hr in the dark. Biofilms were imaged using an LSM-700 confocal microscope with a 63x objective and analyzed using ZEN software. Scale bars are shown in white in each panel and represent 5 µm.

**Figure 5.**
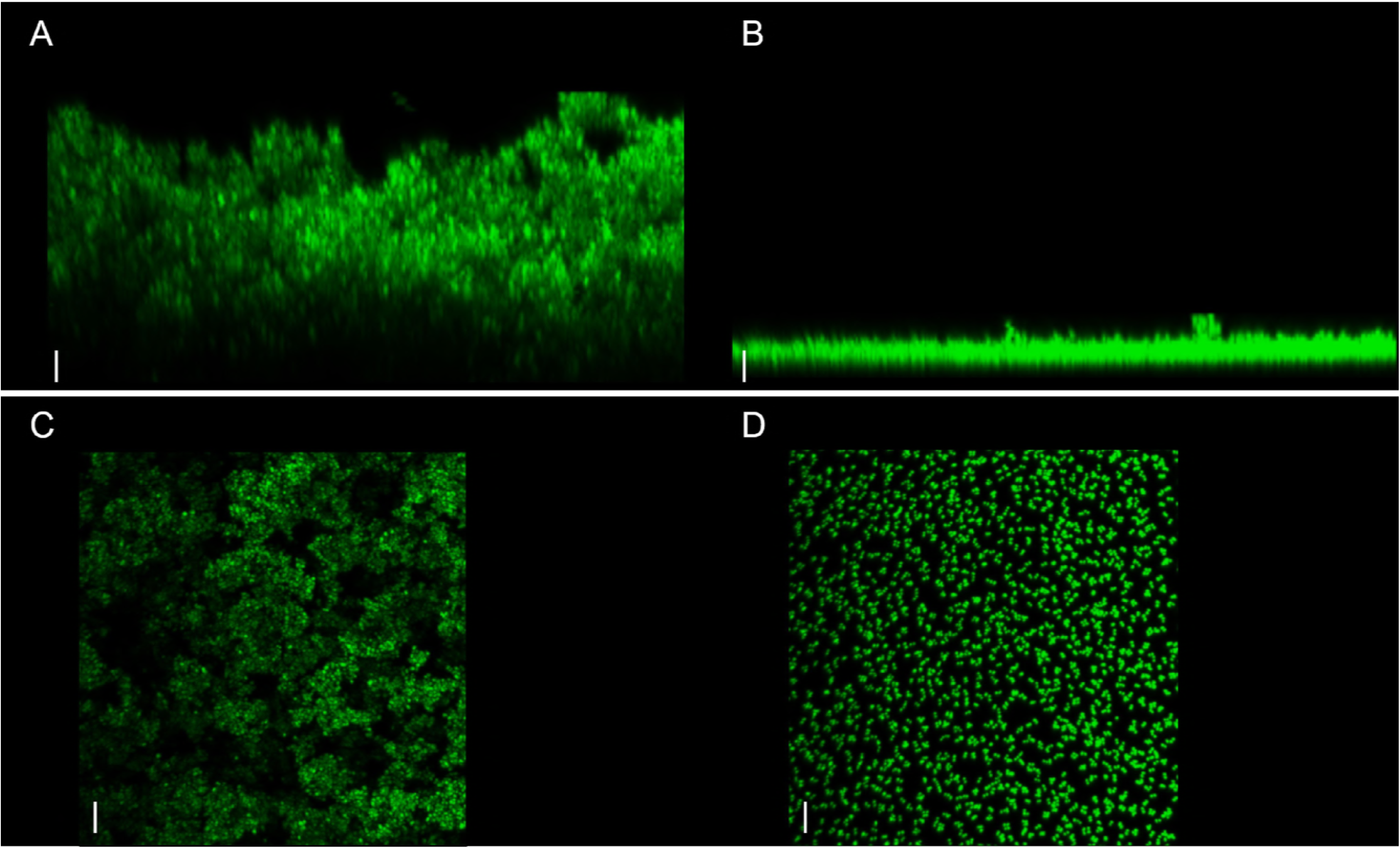
Effect of sublethal concentration of sodium nitrite on *S*. *aureus* biofilms by CSLM. Images show depth (side view) and top view of biofilms formed by wild-type JE2 in acidified TSB media supplemented with 0.75% glucose (aTSB-G) (A and C), and TSB-G media and 20mM sodium nitrite (B and D). Biofilms were formed on the bottom of 6-well polystyrene plates and stained with 2 μM Syto9 for 1 hr in the dark. Biofilms were imaged using an LSM-700 confocal microscope with a 63x objective and analyzed using ZEN software. Scale bars are shown in white in each panel and represent 5 µm.

## Discussion

A number of studies have implicated LMW thiols in biofilm formation. Mutants defective for GSH production are defective for biofilm formation in *P. aeruginosa* (57) and a mutant defective in MSH production is defective for biofilm formation in *Mycobacterium smegmatis* (56). Moreover, the importance of thiols may not be limited to bacterial biofilms. For example, the production of GSH is upregulated in the early stages of biofilm formation by the opportunistic fungal pathogen *Candida albicans* (62). Here we report that a *S. aureus* mutant disrupted in the enzyme encoding for the third and last step of BSH biosynthesis, and thus lacking BSH, was significantly impaired in biofilm formation compared to the wildtype. Treatment with diamide, a thiol oxidant, also resulted in impaired biofilm formation. We also treated the BSH mutant with the common reductant, DTT, to see if DTT could rescue the *bshC* mutant phenotype. Our results indicate that instead of complementing the mutant, the application of DTT resulted in less biofilm formation, suggesting that biofilm formation specifically requires BSH. Trivedi *et al*. reported that treatment with DTT resulted in an increase in the formation of biofilms and also an increase in intracellular thiols in *Mycobacterium tuberculosis* (63). The contradictory findings between *M. tuberculosis* biofilm formation in the Trivedi *et al.* study and *S. aureus* biofilm formation in our experiments upon DTT treatment could potentially be explained by the different concentrations of DTT used in the experiments (2-4 mM for *M. tuberculosis* and 200 mM for *S. aureus*), whereby high DTT concentrations would likely result in oxidative stress instead of reductive stress. Our results support previous studies that indicate that other reducing agents, such as β-mercaptoethanol and cysteine inhibit *S. aureus* biofilm formation, although they were found not to affect the initial adhesion step of biofilm formation (64).

Biofilm formation may be connected to the interaction of LMW thiols and NO. It is likely that NO interacts with BSH during *S. aureus* biofilm formation as our results indicate that the putative BSNOR mutant, which shows a decrease in biofilm formation, also has reduced BSH levels after treatment with acidified sodium nitrite. It is known that NO reacts with GSH to form GSNO, which may serve as a reservoir for NO or conversely sequester GSH so that it is unavailable to the cell (65). NO can *S*-nitrosylate proteins directly through a radical-mediated pathway or indirectly via higher oxides of NO, and may compete with *S*-glutathionylation of exposed cysteines (66). Moreover, GSH can react with protein *S*-nitrosothiols leading to either denitrosylaton or *S*-glutathionylation (67). Schlag *et al*., 2007 demonstrated nitrite dependent inhibition of biofilm formation in *S. aureus* with repression of the icaADBC gene cluster, mediated by IcaR (53), although acidified sodium nitrite was not used in these experiments. Genes involved in DNA repair, reactive oxygen intermediates (ROI), RNS detoxification, and iron homeostasis were induced, and preformed biofilms could be eradicated by the addition of nitrite (53). Major *et al* further demonstrated that acidified sodium nitrite could inhibit biofilm formation and kill planktonic cultures grown aerobically and anaerobically in *S. aureus, P. aeruginosa* and *Burkholderia cepacia* (68). Our results also show that acidified sodium nitrite causes an inhibition of biofilm formation.

Since levels of *S*-nitrosothiols are modulated by *S*-nitrosothiol reductase, which reduces the GSNO to ammonia or other reduced nitrogen species (58), we measured biofilm formation in a putative BSNOR mutant and found that this mutant is defective in biofilm formation. However, the BSNOR mutant was not impaired in growth on formaldehyde or nitrosative stress. Other mutants such as the MscR (dual function *S*-nitrosomycothiol reductase and formaldehyde dehydrogenase) mutant in mycobacteria are impaired in biofilm formation and sensitive to these stressors (56). It has been noted that *S. aureus* is particularly resistant to NO and nitrosative stress as compared to other bacterial species (31). Additionally, a number of *S. aureus* genes have been annotated as aldehyde dehydrogenases, such as *adhA*, which is highly upregulated upon formaldehyde and methylglyoxal treatment (69). It has been suggested that *adhA* functions in the LMW thiol-dependent detoxification of aldehydes, but the gene product has not yet been validated as a BSNOR. Nevertheless, it is possible that there is redundancy in protection against nitrosative and formaldehyde stress in *S. aureus* that leads to a lack of growth phenotypes under these stressors for the BSNOR mutant.

We also assessed the gene expression of BSNOR in wildtype under nitrosative stress and observed no differences in expression levels in the presence or absence of the nitrosative stress. In *Vibrio cholerae*, GSNO reductase activity is regulated post-translationally by *S-*nitrosylation and this activity can be reversed by DTT (70). *S*-nitrosylation of GSNOR1 leading to inhibition of activity has also been documented in plants (71). In addition, in *Neisseria meningitidis, S*- glutathionylation of the cys54 of the thioesterase catalyzing the downstream step from S-formylglutathione hydrolase has been shown to decrease the activity of the enzyme (72). Together, these observations suggest that BSNOR may also be regulated post-translationally, possibly through *S*-bacillithiolation of the corresponding esterase in *S. aureus*, which would result in increased formaldehyde levels and less biofilm formation as seen in *N. meningitidis*. Interestingly, *bshC* expression is decreased under sodium nitrite and hydrogen peroxide and this likely results in a concomitant decrease in *S*-bacillithiolation in *S. aureus* under these stressors.

Recently Gupta *et al.* identified a transcriptional regulator, BifR, which is a member of the MarR protein family in *Burkholderia thailandensis* (73). Under oxidizing conditions, BifR forms a disulfide linked dimer of dimers, which affects its ability to bind to promoters of several genes. A mutant disrupted in BifR had enhanced biofilm formation (73). A similar redox switch controlling biofilm formation may be present in *S. aureus.* In the *bshC* and the BSNOR mutant strains, one would predict that treatment with sodium nitrite would lead to a decrease in BSH and a more oxidized cytoplasm, which would facilitate disulfide formation of a transcriptional regulator, consequently affecting expression of genes involved in biofilm formation. Additionally, in the BSNOR mutant, the BSNO would not be reduced and BSH not released, causing oxidative stress further affecting the regulation of biofilm formation. Whether this occurs in *S. aureus* needs further study.

The data presented here indicates that the bacillithiol pathway likely interacts with NO and together they play key roles in biofilm formation in *S. aureus*. Further characterization of the mechanistic details of this interaction and a search for a transcriptional regulator or other proteins that link these pathways will further our understanding of oxidative and nitrosative stress mechanisms in biofilm formation in *S. aureus*, and may lead to the development of novel targets for therapies to eliminate biofilms in the clinic.

## Materials and Methods

### Strains and culture conditions

*S. aureus* USA300 (JE2) wild-type cultures were grown from glycerol stocks on TSA (Tryptic Soy Agar) (BD211825) plates and incubated at 37°C overnight. Antibiotics were added when appropriate (erythromycin at 10 µg/ml for mutants *bshC (NE230)* and *NE122* disrupted in SAUSA300_0055, a putative *S*-nitrosobacillithiol reductase. These transposon mutants were provided by the Network on Antimicrobial Resistance in *Staphylococcus aureus* (NARSA) for distribution by BEI Resources, NIAID, NIH. For all cultures, a single colony was inoculated in TSB (Tryptic Soy Broth) (BD211825) liquid media and grown at 37°C, shaking at 250 rpm, for 12 hours. All biofilms, except experiments concerning sodium nitrite, were grown in TSB media supplemented with 0.75% glucose. For experiments with sodium nitrite, biofilms were grown in acidified TSB media (pH 5.5) supplemented with 0.75% glucose. Biofilms were grown as follows: cell cultures grown for 12 hours were diluted to a final OD_600_ of 0.2 and added to all wells excluding blank (no cells added) control wells. The plate was incubated for 60 minutes at 37°C under static conditions, wells were washed with PBS and 100 µl of media was added. The plate was incubated for 24 hours at 37°C under static conditions. The media was aspirated, and the biofilm was measured by OD_600_. Experimental data for each concentration was obtained by subtracting the OD_600_ readings of the average blank well of each concentration from each corresponding experimental well.

### Minimum inhibitory concentrations (MICs)

For planktonic MICs, DTT (Fisher scientific BP17225), diamide (MP Biomedicals 0210152705) and sodium nitrite (VWR AA 14244-22) were serially diluted two-fold from 1M to 61 µM in 100 µl TSB. Acidified TSB media (pH 5.5) was used for sodium nitrite. Cell cultures grown for 12 hours, diluted to a final OD_600_ of 0.005 and added to all wells excluding blank (no cells added) control wells. The 384-well plate (Thermo 242765) was incubated at 37°C for 24 hours with no shaking. Eleven replicates were performed for each concentration and the MIC assay was performed in replicate. The lowest concentration at which cell turbidity was not visible was determined to be the MIC.

For biofilm MIC, DTT, diamide and sodium nitrite were serially diluted two-fold from 1M to 61.04 µM in 100 µl TSB, supplemented with 0.75% glucose or 100 µl acidified TSB (pH 5.5), supplemented with 0.75% glucose (for sodium nitrite). Cell cultures grown for 12 hours were diluted to a final OD_600_ of 0.2 and added to all wells excluding blank (no cells added) control wells. The plate was incubated with no shaking at 37°C for 60 minutes, wells were washed with PBS and 100 µl of media with DTT, diamide or sodium nitrite (serially diluted twofold from 1M to 61 µM) was added. The plate was incubated for 24 hours at 37°C under static conditions. The media was aspirated, and the biofilm was measured by OD_600_. Experimental data for each concentration was obtained by subtracting the OD_600_ readings of average blank well of each concentration from each corresponding experimental well. The blank subtracted OD_600_ values of each experimental well (10 replicates per concentration) was divided by the blank subtracted OD_600_ value of control well (no DTT or diamide added) to obtain normalized values. Reported data represents mean normalized value and standard deviation.

### Confocal scanning laser microscopy (CSLM) biofilm visualization

Cell cultures grown for 12 hours, were diluted to a final OD_600_ of 0.2 in 4 ml in a 6-well plate (Falcon 351146). For biofilm formation without added drugs, JE2, BshC and BsnoR, cells were added to TSB, supplemented with 0.75% glucose. For biofilms formed in the presence of DTT or diamide, 1.95 mM diamide or 150 mM DTT in TSB supplemented with 0.75% glucose was used. For sodium nitrite experiments, acidified TSB (pH 5.5) supplemented with 0.75% glucose and 20 mM sodium nitrite or no sodium nitrite was used. The 6-well plates were incubated for 60 min at 37°C, wells were washed with PBS and 4 ml of fresh media was added to each well. The plates were incubated for 24 hrs at 37°C with no shaking. Biofilms were stained with 5 µM Syto9 nucleic acid stain (Thermo Fisher S34854) for 1 hour at 37°C in the dark. Medium containing the stain was removed, 4 ml of PBS was added, and biofilms were imaged using an LSM 700 confocal microscope with a 63x objective. Images were analyzed, and biofilm thicknesses were measured using ZEN software (Carl Zeiss).

### Analysis of LMW thiol levels after treatment with sodium nitrite

*S. aureus* cultures were grown overnight in tryptic soy broth (TSB) before being diluted in 50 mL acidified TSB (pH 5.5) to either OD_600_ 0.2, or 0.1 and incubated at 37 **°**C. Quadruplicates were treated with 20 mM sodium nitrite once cultures reached OD_600_ 0.5 for 30 minutes at 37 **°**C. Immediately after treatment, 10 µL of culture was serial diluted in PBS by 10^-5^, 10^-6^, and 10^-7^, and 50 µL was plated on tryptic soy agar and incubated at 37 **°**C to determine if significant killing occurred. The remaining culture was pelleted by centrifugation at 4,000 RPM for 10 minutes at 4**°**C and the supernatants were removed. Pellets were stored at −80 **°**C overnight. LMW thiol analysis was performed on pellets as previously described (40).

### Quantative real-time pCR (qPCR)

Expression levels of *katA, BSNOR* and *bshC* treated with 20 mM sodium nitrite for 5 min or 30 min in the *S. aureus* wildtype strain were measured by qPCR. Samples were grown in acidified (pH 5.5) tryptic soy broth (TSB) until they reached an OD_600_ of 0.5 before treatment. Relative gene expression was calculated using the −2^ΔΔCT^ method (ΔC_t_ = C_t sample_ - C_t control_) and reported as fold change in gene expression of each sample normalized to the *gyrB* housekeeping gene relative to the untreated culture. A Mann-Whitney U test was performed to determine significance p<0.05.

## Acknowledgements

We thank all members of the Rawat and Nobile labs as well as Aaron Hernday for insightful discussions and comments on the manuscript. The confocal scanning laser microscopy images in this work were collected, in part, with the use of a confocal microscope acquired through the National Science Foundation (NSF) MRI award DMR-1625733. This work was supported by the National Institutes of Health (NIH) National Institute of General Medical Sciences (NIGMS) award R35GM124594 to CJN, and by the NIH NIGMS award SC3GM100855 and National Science Foundation (NSF) award MCB 1244611 to MR. The funders had no role in study design, data collection and interpretation, or the decision to submit the work for publication.

## Conflict of Interest

Clarissa J. Nobile is a cofounder of BioSynesis, Inc., a company developing inhibitors and diagnostics of biofilm formation, and Megha Gulati is a consultant of BioSynesis, Inc.

---

**Supplemental Figure 1: *S*. *aureus* cultures grown in nutrient rich media.** Overnight cultures were grown in tryptic soy broth (TSB) at 37**°**C while shaking before being diluted to OD_600_ 0.05 in 200 uL in a 96-well plate with indicated treatments. *S. aureus* wildtype (green), *bshC*_-_ (blue), and *BSNOR-* (purple) were incubated at 37**°**C for 24 hrs with reads every hour. *S. aureus* was either non-treated, treated with GSNO at 1 mM or 5 mM, treated with formaldehyde at 0.25 mM, 0.75 mM, or 1.5 mM, treated with acidified sodium nitrite pH 5.5 (20 mM), or treated with methylglyoxal at 0.75 mM or 1.5 mM.

